## Supporting figures and legends for "R-loops acted on by RNase H1 are a determinant of chromosome length-associated DNA replication timing and genome stability in *Leishmania*"

#### Supplementary figures and legends

**Supplementary Figure S1. Detection of R-loops around RNA Pol II transcribed regions in *L. major*.** Snapshot of DRIP-seq at the indicated genomic regions; from top to bottom: track 1 and 2 (green), R-loop enriched regions relative to input material; -RNase H and +RNase H indicate mock or treatment with recombinant RNase HI prior to immunoprecipitation, respectively; R-loop peaks are indicated as purple bars bellow track 1; track 3 (dark red), MNase-seq data; track 4 (blue), mapping of G quadruplex structures (G4s); black arrows at the bottom of each panel indicate annotated coding sequences (CDSs).

**Supplementary Figure S2. Detection of R-loops around RNA Pol I and Pol III transcribed regions in *L. major*. A)** and **B)** Snapshot of DRIP-seq at Pol I and Pol III transcribed regions, respectively; from top to bottom: track 1 and 2 (green), R-loop enriched regions relative to input material; -RNase H and +RNase H indicate mock or treatment with recombinant RNase HI prior to immunoprecipitation, respectively; R-loop peaks are indicated as purple bars bellow track 1; track 3 (dark red), MNase-seq data; track 4 (blue), mapping of G quadruplex structures (G4s); black arrows at the bottom of each panel indicate annotated coding sequences (CDSs).

**Supplementary Figure S3. Genome-wide R-loop distribution. A)** Colourmap showing DRIP-seq in all 36 chromosomes of wild type (*WT*) cells; chromosomes are ordered by size; enrichment was calculated either as the enrichment of immunoprecipitated material relative to input (IP/Input; two panels at the left) or enrichment of immunoprecipitated material from mock relative to immunoprecipitated material from pre-treatment with recombinant RNase HI (IP; panel at right); -RNase H and +RNase H indicate mock or treatment with recombinant RNase HI prior to immunoprecipitation, respectively. **B)** Snapshot showing DRIP-seq in the indicated chromosomes; chromosome length is indicated in parenthesis; position and orientation of polycistronic transcription units (PTUs) is indicated for each chromosome.

**Supplementary Figure S4. Analysis of correlation between sequence content and chromosome length. A), B), C)** and **D)** Colourmaps showing density of the indicated genome features; chromosomes are ordered by size. **E), F), G)** and **H)** Linear regression analysis between chromosomes size and the indicated genomic features; R and *P* values are indicated at the top of each panel; shaded areas indicate 95% confidence interval.

**Supplementary Figure S5. Using CRISPR/Cas9 to generate an *RNaseH1-HA$^{Flox}$* cell line. A)** CRISPR-Cas9 was used to flank the *RNase H1* ORF with *LoxP* sites and fuse it with HA tag (*RNase H1-HA$^{Flox}$*); *a, b, c* and *d* indicate the annealing position of primers used in B. **B)** PCR analysis of genomic DNA extracted from the DiCre/Cas9/T7-expressing cell line and the *RNase H1-HA$^{Flox}$* cell line; primer annealing positions are shown in A. **C)** Growth curve of *RNase H1-HA$^{Flox}$* cell line (grey) as compared to wildtype (WT) cells (black); cells were seeded at $2x10^5$ cells.$mL^{-1}$ at day 0; cell density was assessed every 24 h, and error bars depict standard error of the mean (SEM). **D)** PCR analysis of genomic DNA extracted from WT cells and an *RNase H1 KO* clonal cell line; amplification of the *Rad51-3* gene was used as loading control.

**Supplementary Figure S6**. **Workflow for KO induction in the *RNase H1-HA$^{Flox}$* cell cline**. Exponentially growing cells were seeded in medium with (+RAP) or without (-RAP) rapamycin; every 4 - 5 days of cultivation cells were re-seeded; all the experiments reported here were performed in cells subjected to this induction protocol.

**Supplementary Figure S7**. **Demonstration of *RNase H1* KO induction by whole genome sequencing**. Whole genome DNA sequencing (top 3 panels) and whole genome RNA sequencing (bottom 3 panels)**;** RNase H1 gene position is indicated by a dashed red line.

**Supplementary Figure S8**. Quantification of R-loop levels as detected via immunofluorescence using S6.9 antibody upon the indicated conditions; each treatment was performed for 6 hours.

**Supplementary Figure S9**. **Cell cycle progression analysis and quantification of S phase cells upon *RNase H1* KO. A)** Exponentially growing cells were left untreated (N.T.) or treated for 8 hours with 5 mM hydroxyurea (HU) and then re-seeded in HU-free medium; cells were collected at the indicated time points after HU removal, fixed, stained with Propidium Iodide (P.I.) and analysed by FACS; 2C and 4C indicate one DNA content (G1) and double DNA content (G2/M), respectively. **B)** Exponentially growing cells were pulsed with BrdU for 30 minutes; BrdU fluorescence was detected under denaturing conditions using anti-BrdU antibody; 30,000 cells were analysed per condition; dashed red lines indicate the BrdU-positive population, i.e. cells in S phase; inset numbers indicate percentage of cells in S phase.

**Supplementary Figure S10. Mutation signature analysis. A)** Representation of the proportion of transition (Ts) and transversion (Tv) SNPs at the indicated conditions. **A)** SNP plus 20 flanking nucleotides was used for pLogo fold enrichment analysis; SNPs were centred in each plot (vertical dotted black line); overrepresented nucleotides are above, and underrepresented are below the horizontal dotted black line; horizontal red lines indicate threshold of significant enrichment ($p < 0.05$); font size indicates the enrichment magnitude.

### Supplementary Figure S1

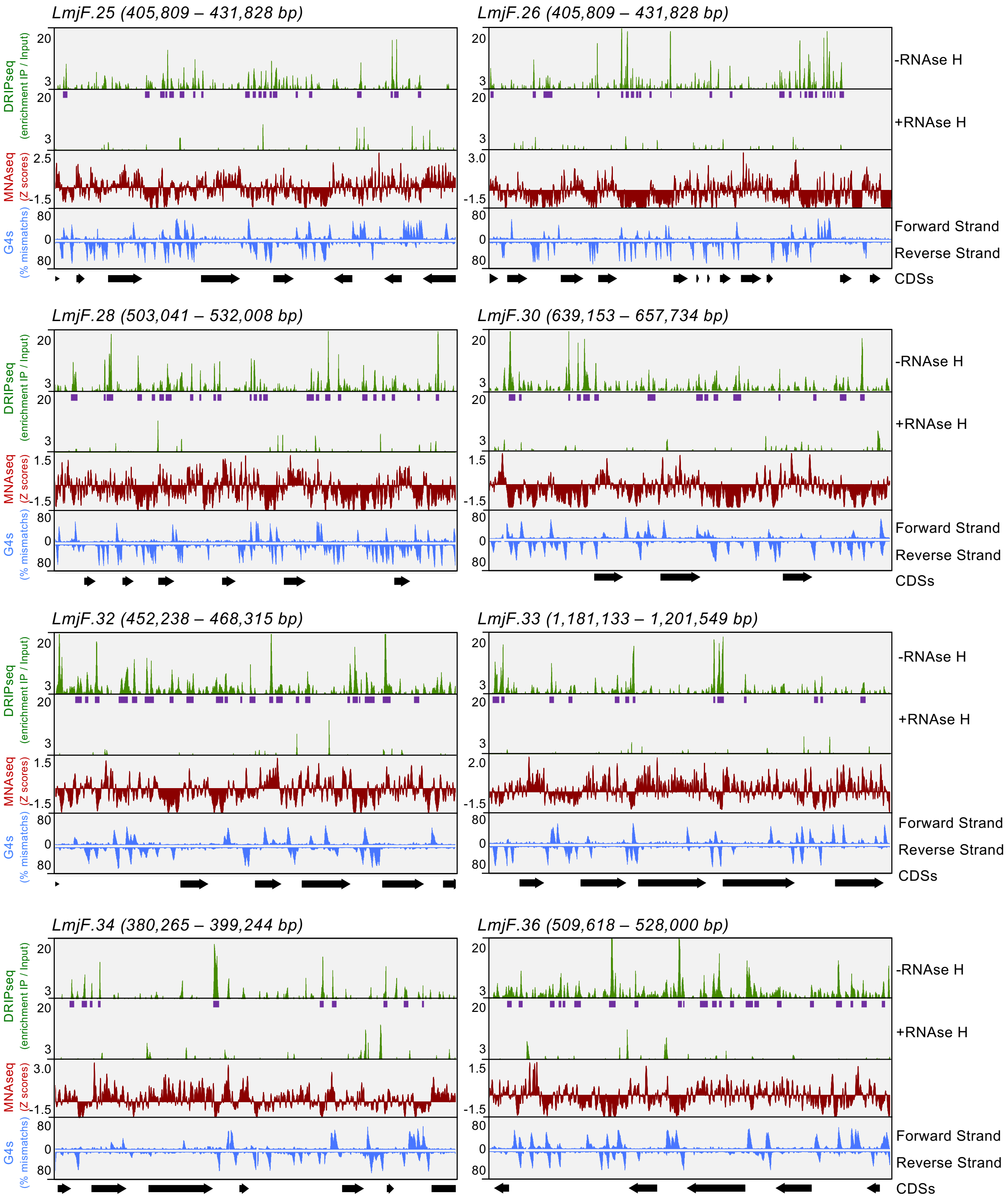

### Supplementary Figure S2

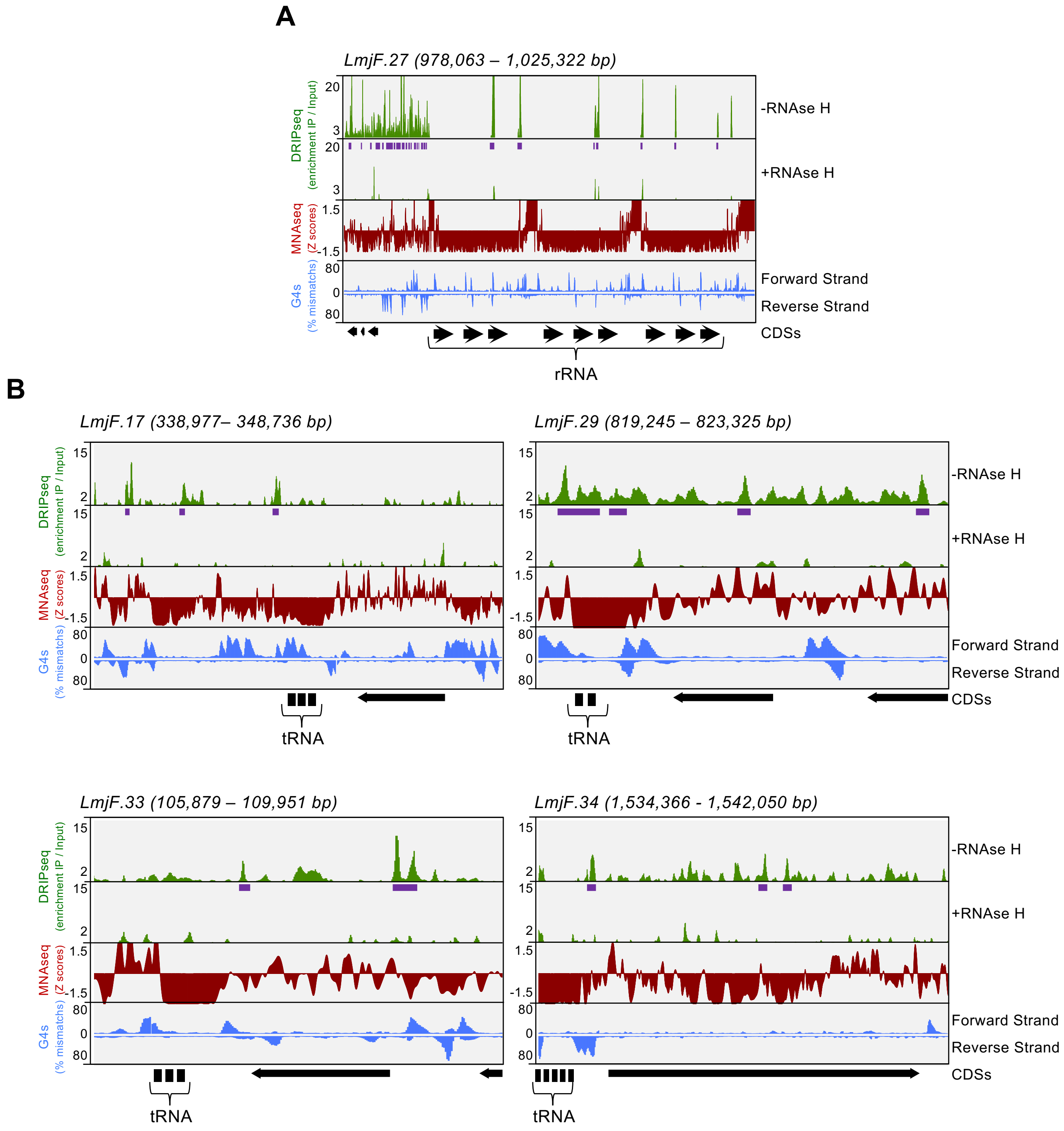

### Supplementary Figure S3

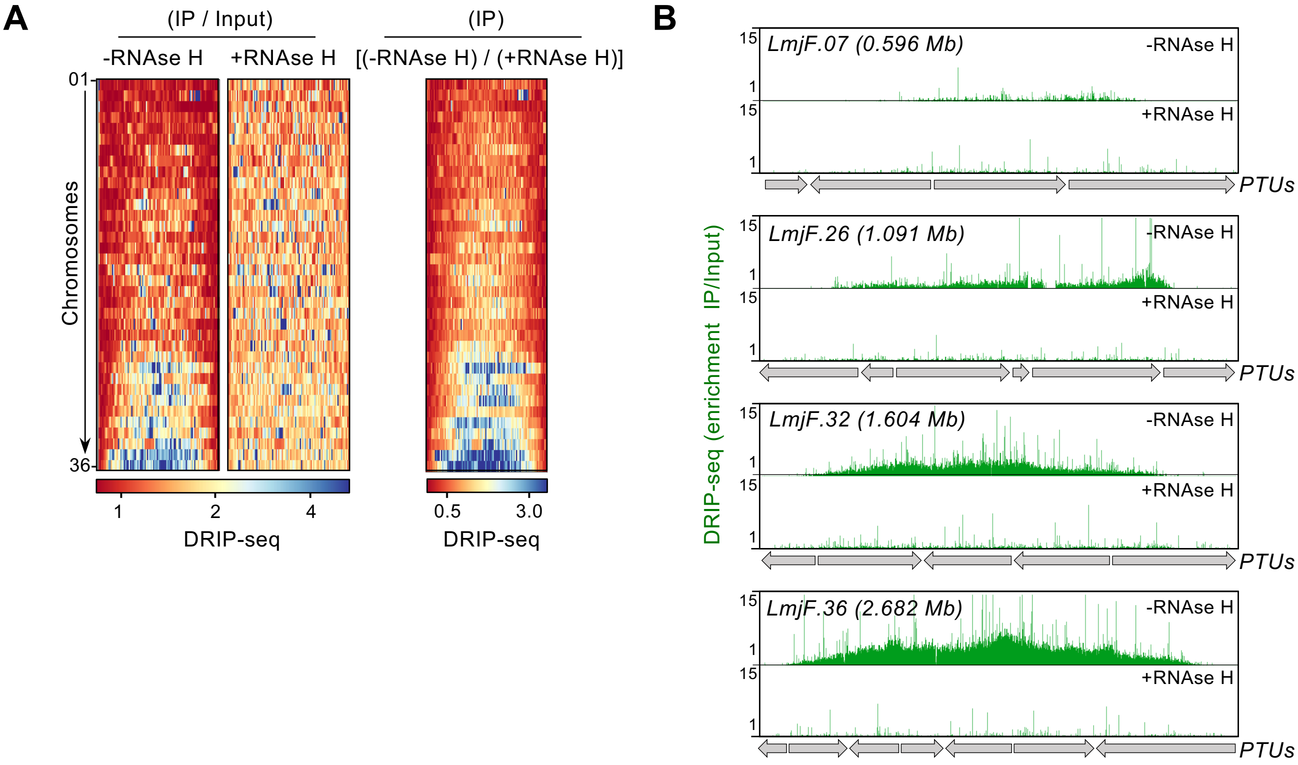

### Supplementary Figure S4

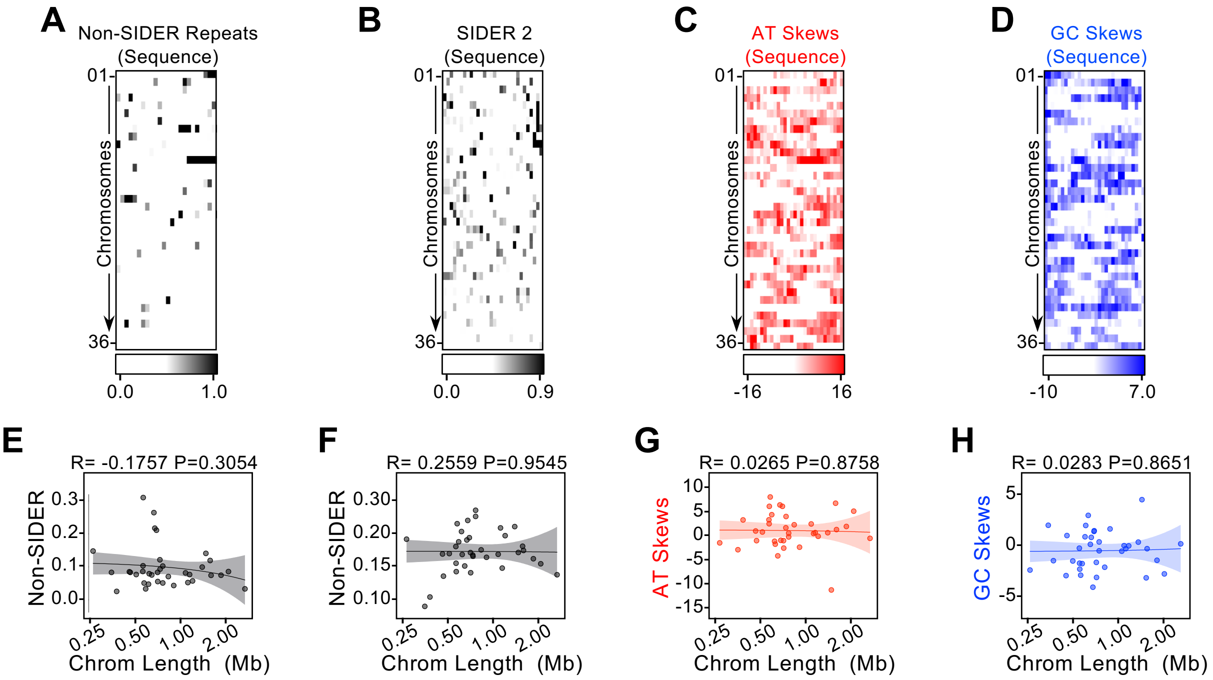

### Supplementary Figure S5

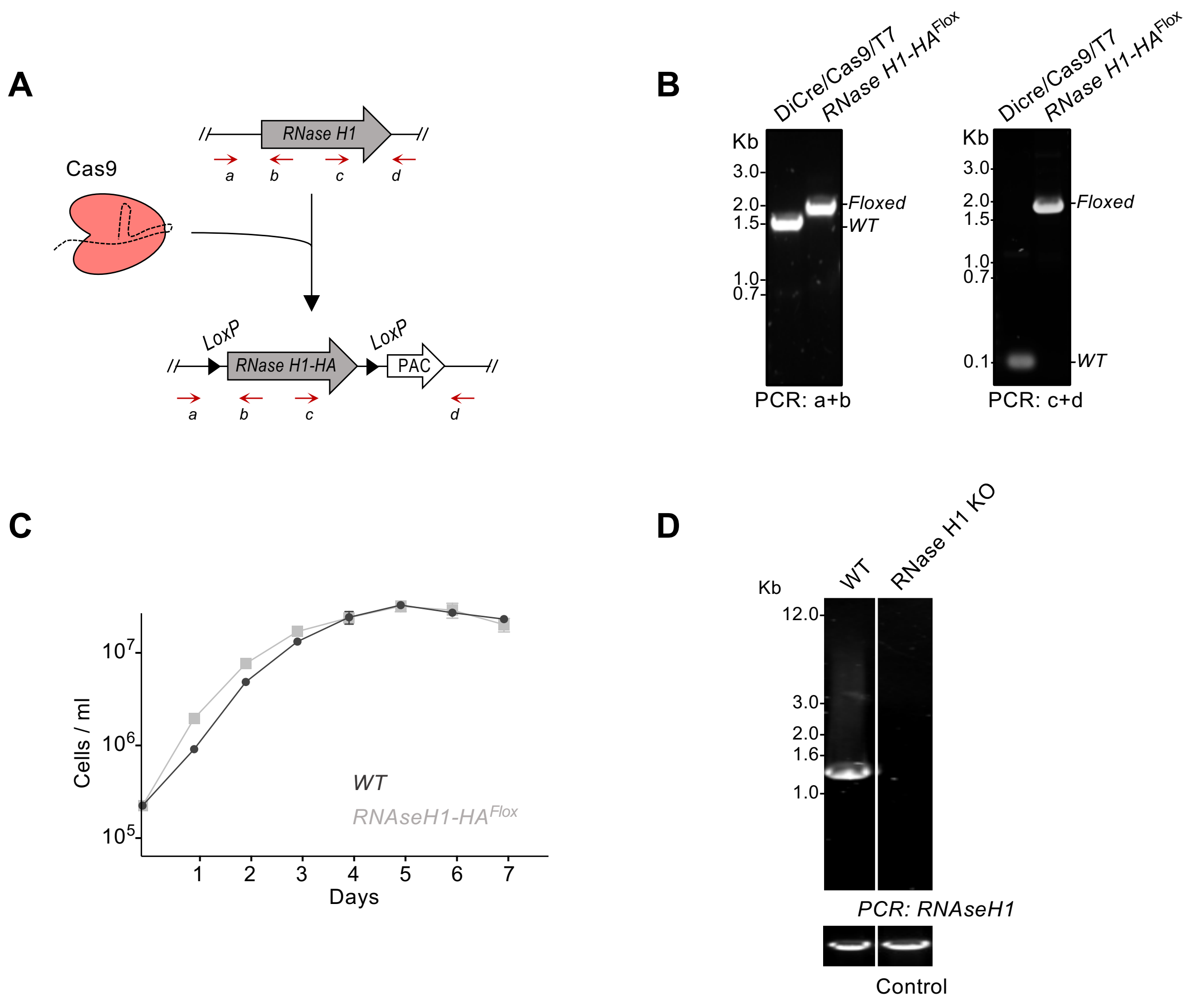

### Supplementary Figure S6

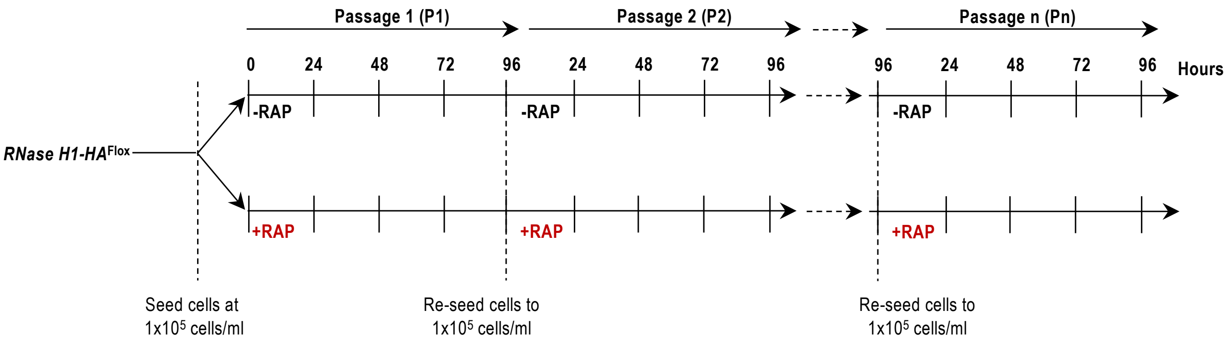

### Supplementary Figure S7

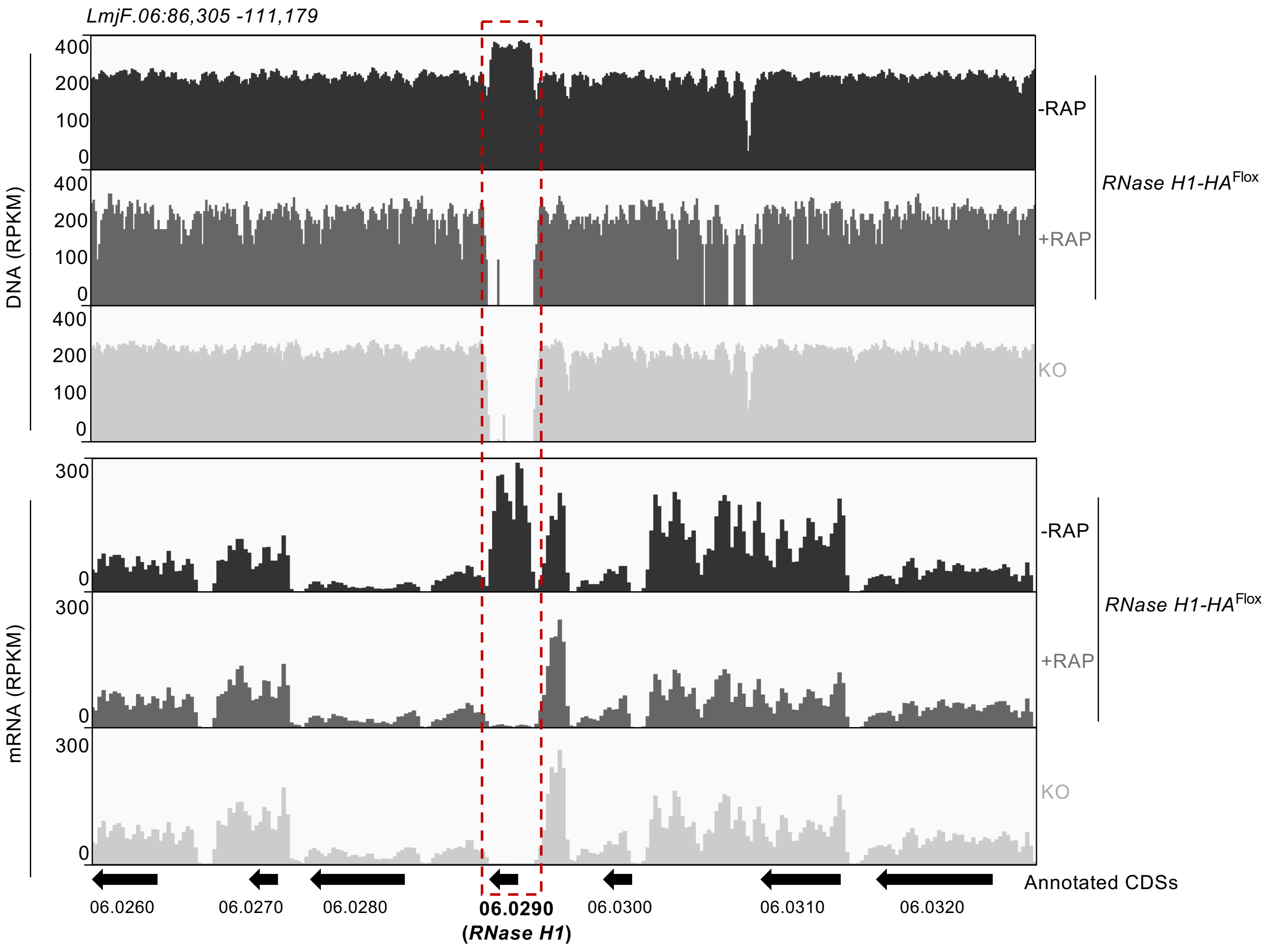

### Supplementary Figure S8

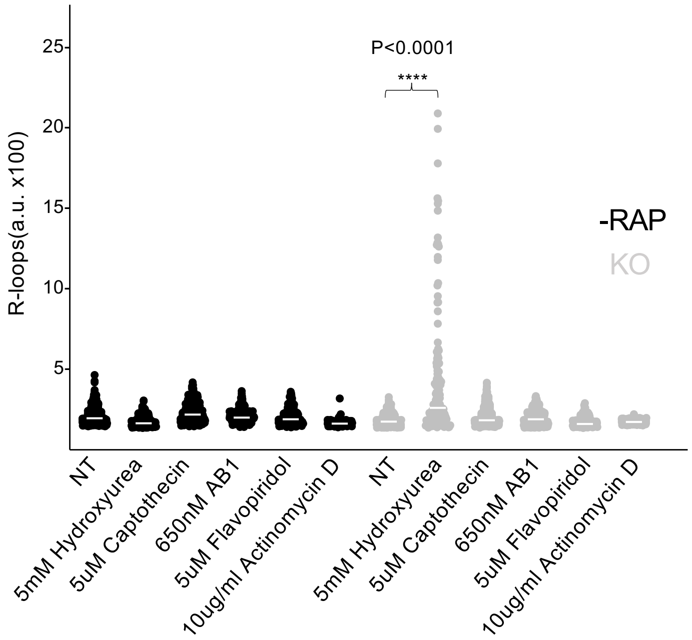

### Supplementary Figure S9

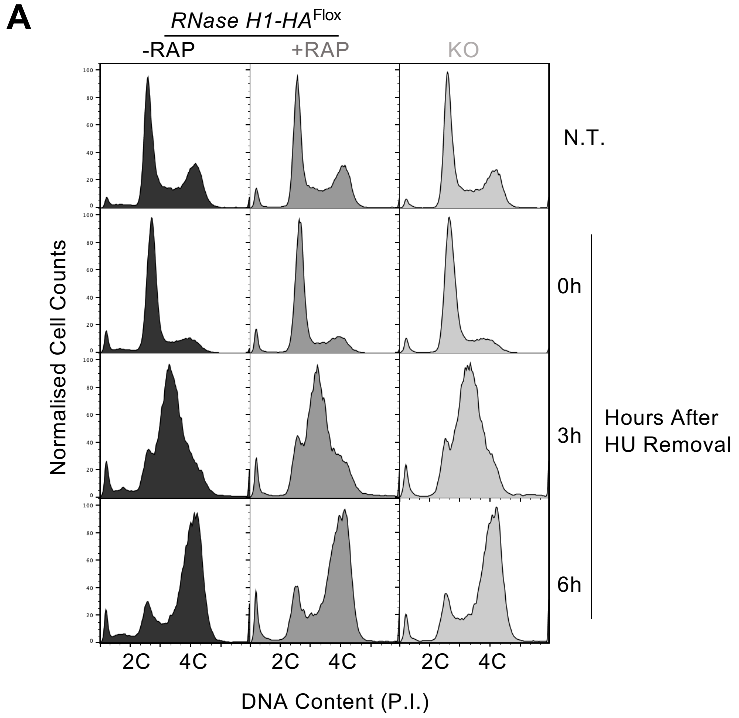


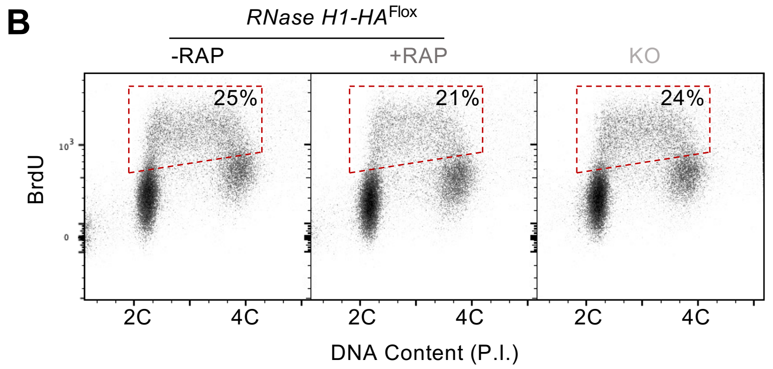

#### Supplementary Figure S10

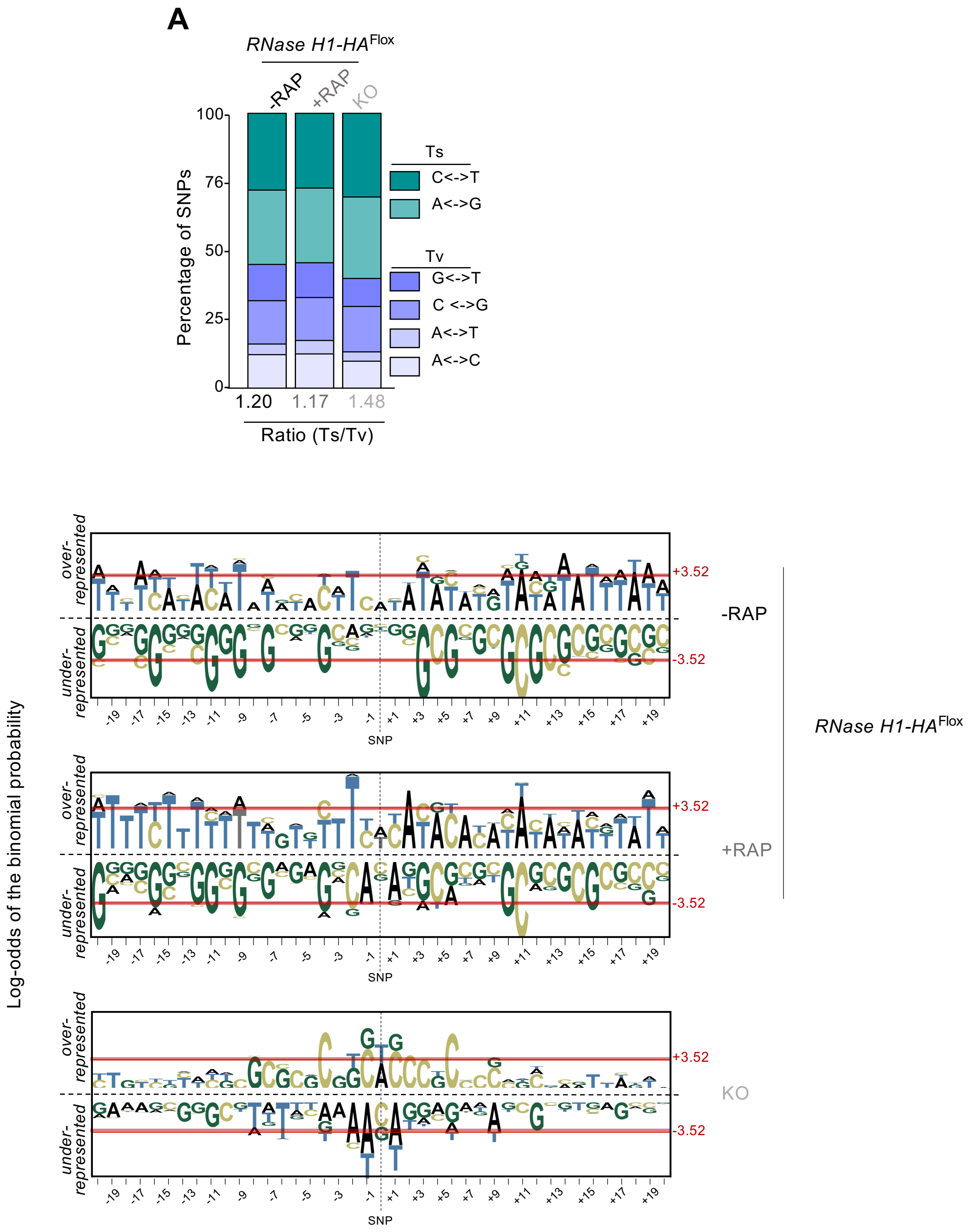
